# Matrix metalloproteinase-3 promotes arteriovenous fistula failure by regulating FAK-AKT signaling

**DOI:** 10.1101/2025.08.27.672378

**Authors:** Yangzhouyun Xie, Weichang Zhang, Carly Thaxton, Ying Jin, Bogdan Yatsula, Hualong Bai, Sean Davis, Tobias Exsted, Alan Dardik, Raul J Guzman, Yujun Cai

**Author notes:** Corresponding authors: Yujun Cai, PhD Division of Vascular Surgery and Endovascular Therapy Department of Surgery Yale School of Medicine 310 Cedar St. BML226 New Haven, CT 06510 Raul J. Guzman, MD Division of Vascular Surgery and Endovascular Therapy Department of Surgery Yale University School of Medicine New Haven, CT 06510 Alan Dardik, MD, PhD. Vascular Biology and Therapeutics Program Yale School of Medicine 10 Amistad Street, Room 437 New Haven, CT, 06510.

## Abstract

**Objective-** Surgically created upper extremity arteriovenous fistulae (AVF) are the preferred vascular access for patients requiring dialysis. It is estimated, however, that 50% of AVF fail within one year due to aggressive neointimal hyperplasia, which significantly increases morbidity and mortality. Matrix metalloproteinase-3 (MMP-3), also known as stromelysin-1, is a member of the metalloproteinase family that plays a critical role in the pathogenesis of many human disorders by degrading extracellular matrix and regulating molecular signaling pathways. The role of MMP- 3 in AVF neointimal failure has not been explored. **Approach and Results**- We observed that MMP-3 was induced in a time-dependent fashion by fetal bovine serum (FBS) and the growth factor PDGF-BB in cultured venous SMC. MMP-3 was also highly expressed in the neointimal SMCs of the outflow veins and the juxta-anastomotic area in an AVF mouse model, as well as in human AVF specimens. Knockdown of MMP-3 significantly suppressed venous SMC proliferation, whereas overexpression of MMP-3 facilitated cell growth *in vitro*. Importantly, deficiency of global and SMC-specific MMP-3 significantly reduced neointimal hyperplasia and improved patency after AVF creation. Mechanistic studies showed that MMP-3-mediated SMC proliferation and AVF neointimal hyperplasia were regulated via the FAK-AKT signaling pathway. **Conclusions**- These data suggest that MMP-3 is a key mediator of AVF neointimal failure. Targeting local MMP-3 activity may be a novel therapeutic strategy to prevent AVF neointimal failure and improve outcomes in patients requiring hemodialysis.

## Introduction

Vascular access is required for hemodialysis in patients with end-stage kidney disease (ESKD). There are three types of vascular access available: arteriovenous fistulae (AVF), arteriovenous grafts, and catheters. Among these, AVF is the preferred vascular option due to increased long-term patency and utilization.^1^ However, AVF can fail due to stenosis in the juxta-anastomotic area and the outflow vein.^2–4^ It has been estimated that the one-year patency rate of AVF is only 50%.^5–7^ The leading cause of AVF failure is neointimal hyperplasia, characterized mainly by vascular smooth muscle cells (SMC) proliferation and extracellular matrix (ECM) deposition.^8–11^ This results in narrowing of the venous outflow lumen and subsequent dysfunction during dialysis or occlusion. Understanding molecular mechanisms related to SMC proliferation and neointima formation will help develop new strategies aimed at improving AVF patency and longevity.

Matrix metalloproteinases (MMP) are a superfamily containing a large number of members that play critical roles in many pathophysiological processes, including wound healing, vascular remodeling, and angiogenesis.^12–14^ In addition to degrading extracellular matrix, MMPs are involved in regulating intracellular signaling pathways such as cell migration, proliferation, and apoptosis.^15–18^ MMP-3, also called stromelysin-1, is a member of the MMP family that plays important roles in various human disorders.^19, 20^ MMP-3 is implicated in many pathological vascular disorders, including remodeling, atherosclerosis, and abdominal aortic aneurysms.^21, 22^ Recently, we have shown that MMP-3 in SMC is an important mediator of arterial calcification.^23^ We have also observed that MMP-3 is largely induced (>180-fold) in response to AVF creation *in vivo*,^10, 24^ suggesting its involvement in AVF vascular remodeling.

The protein tyrosine kinase focal adhesion kinase (FAK) plays critical roles in regulating cell growth and survival.^25–27^ FAK has been evaluated as a therapeutic target for cancer therapy, and several FAK inhibitors are currently in phase II clinical trials for resistant non-small cell lung cancer. ^28–30^ Recent studies have also demonstrated that FAK is implicated in many vascular diseases including restenosis, atherosclerosis, and pulmonary hypertension.^26, 31, 32^ FAK is primarily located in SMC nuclei in healthy vessels, however, in response to vascular injury, it can translocate from the nuclei to the cytoplasm, which ultimately leads to neointimal formation, suggesting the importance of FAK in regulating pathological vascular remodeling.^33–35^ FAK activation can induce PI3K/AKT signaling that contributes to cell survival and proliferation, suggesting that FAK-AKT signaling could be critical for pathological vascular remodeling.^36, 37^

In this study, we sought to explore the role of MMP-3 in AVF failure. Using a surgically created AVF model in mice, AVF specimens from patients with ESKD, MMP-3 transgenic mice, and *in vitro* mechanistic studies in SMCs, we demonstrate that MMP-3 is a key mediator in AVF neointimal hyperplasia and failure, and the underlying mechanisms primarily involve regulation of FAK-AKT signaling.

## Methods

### Reagents

MMP-3 inhibitor II, tamoxifen, and corn oil were obtained from Millipore Sigma (St. Louis, MO). Dulbecco’s Modified Eagle’s Medium (DMEM), and fetal bovine serum (FBS) were purchased from ThermoFisher Scientific (Waltham, MA). VS-4718, RNase A and Propidium Iodide were purchased from MedChem Express (Monmouth Junction, NJ).

### Mice

Animal care and procedures were conducted in accordance with animal protocols (IACUC 2022- 2081, 2022-20279) that were approved by the University Animal Care and Use Committee at Yale University and National Institutes of Health (NIH) guidelines. Global MMP-3 knockout mice were previously provided by Lynn Matrisian, PhD when at Vanderbilt University. This mouse line has undergone at least 9 generations of backcrossing with C57BL/6 mice. MMP-3 floxed mice and SMC-specific MMP-3-deficient mice were generated as we previously described.^23^ Briefly, MMP- 3 floxed mice were created using CRISPR/Cas9 technology and subsequently crossed with SMC-specific Cre mice SMMHC-CreER^T2^ (JAX, strain #: 019079) to generate SMMHC-Cre^+^/MMP-3- flox^+/+^ mice. To induce SMC-specific MMP-3 deficiency, SMMHC-Cre^+^/MMP-3-flox^+/+^ mice received intraperitoneal injections of 75 mg/kg tamoxifen (TMX) for 5 consecutive days, followed by a 10-day rest period. Animals were anesthetized with 2.5% isoflurane via a vaporizer and euthanized by perfusion with saline. Vessels were directly dissected for Western blotting or animals were further perfused with 10% phosphate-buffered formalin (10% NBF) for immunostaining.

### Mouse aortocaval fistula model

The mouse aortocaval fistula model was used to assess neointimal hyperplasia and failure.^38^ 9- week-old male mice were anesthetized with 2.5% isoflurane via a vaporizer. After buprenorphine SR injection, an abdominal midline incision was performed, and a micro clip was applied to the proximal infrarenal abdominal aorta. A 45-degree curved 25-gauge needle was used to puncture the abdominal aorta, creating an aortocaval fistula by entering the inferior vena cava (IVC). Upon needle withdrawal, the puncture site on the aorta was immediately compressed with surrounding tissue to stop bleeding. The incision was then closed with a 6-0 suture. Successful creation of the AVF was confirmed by increased blood flow and diameter in the IVC as measured by Doppler ultrasound.

### Human specimens

Resected vein segments from human AVF specimens were collected from patients during a second stage basilic vein transposition. Control vein sections not subjected to AVF creation served as controls. We collected de-identified specimens. This study was not considered human subject research as determined and approved by the Yale University Institutional Review Board. All procedures were conducted in accordance with the IRB protocol (1000032195 and 2000038049). Each specimen was divided, with one half snap-frozen in liquid nitrogen for RNA and protein preparation, and the other half fixed in 10% neutral buffered formalin (NBF) for histological and immunostaining.

### Ultrasound

The Ultrasound Vevo770 High-Resolution Imaging System (VisualSonics, Bothell, WA) with probe RMV704 (20–60 MHz) was utilized to monitor the blood velocity, diameter, and patency of the inferior vena cava (IVC) and aorta as described previously.^39, 40^ The waveform in the IVC and the aorta, just cranial to the renal veins, was recorded using the pulse-wave mode. Ultrasound assessments were performed 1 day before and after the creation of the AVF, on days 3, 7, 14, and 21. The patency of the AVF was determined by the presence of turbulent waveforms in the IVC. Blood flow and shear stress were calculated using the following equations: Flow= 60×V×ρχ×r^2^; Shear stress= 4×V×1/r; where V is blood velocity (cm/s), r is the radius (cm), and 1 is blood viscosity, assumed to be constant at 0.035 poise.

### Morphometric Analysis

At the end of the experiment, mice were euthanized and perfused with saline followed by 10% neutral buffered formalin (NBF). The AVF was dissected en bloc, fixed in 10% NBF, and embedded in paraffin, and 5-μm cross-sections were cut. Verhoeff-Van Gieson (VVG) staining was utilized to visualize elastic laminae in the IVC and aorta. To assess wall thickness and neointimal area in the outflow veins, sections were obtained 50-100 μm cranial to the fistula insertion site as described previously.^39, 40^ Wall thickness was determined at eight equidistant points per cross-section and averaged. The intimal plus medial area was calculated as the difference between the external elastic lamina area and the luminal area. For analysis of the juxta-anastomotic area, sections were taken at the largest puncture site of the AVF. This region was defined as the junction where the elastin lamellae disappeared at the confluence of the aorta, IVC, and fistula. Wall thickness measurements in the outflow and the juxta-anastomotic area of the IVC were conducted using ImageJ software.

### Rat and human vein SMC culture

Rat primary vein smooth muscle cells (SMCs) (CellBiologics, RA-6086) and primary human vein smooth muscle cells (CellBiologics, H-6086) were cultured and maintained in a complete smooth muscle cell medium (Cell Biologics, M2268) in a 37°C, 5% CO_2_ humidified incubator. When performing experiments, SMCs (passage 4-8) were cultured in Dulbecco’s Modified Eagle’s Medium (DMEM) (Gibco, 119950065) supplemented with 10% FBS, and 1% of penicillin-streptomycin.

### Mouse aortic SMC isolation and culture

Mouse aortic SMCs were isolated from the thoracic aorta using an enzymatic method.^41^ Briefly, the aorta was dissected and placed in a serum-free DMEM medium on ice. After removal of surrounding fat tissue, the vessel was subjected to adventitia digestion using a buffer containing 161 U/ml Collagenase Type II (Worthington CLS2) and incubated at 37°C for 15 min. Subsequently, the aorta was transferred to serum-free DMEM, the adventitia was peeled off, and endothelial cells were removed by gently scraping with a cotton swab. Next, the aorta was placed in a media digestion buffer containing 402 U/ml Collagenase Type II, cut into pieces, and incubated for 4-6 h. The resulting cell mixture was centrifuged, resuspended in DMEM culture medium supplemented with 20% FBS and 1% penicillin/streptomycin, and cultured in a humidified incubator (37°C, 5% CO_2_). SMCs were confirmed using immunofluorescence staining with anti- SM-α-actin antibody (Dako, M0851). Passage 4-8 cells were used for subsequent experiments.

### Cell proliferation assessment

Sulforhodamine B (SRB) colorimetric assay and BrdU immunofluorescence staining were used to examine SMC proliferation as described previously.^42^ For the SRB assay, rat and human vein SMCs were seeded into 96-well plates and cultured under serum-free conditions for starvation. Subsequently, cells were stimulated with 10% FBS for 48 h. Following stimulation, cells were fixed with 10% trichloroacetic acid for 30 min at 4°C, stained with 4 mg/ml SRB dissolved in 1% acetic acid for 15 minutes, and then solubilized with 10 mM Tris base. Optical density (OD) at 515 nm was measured using a VERSAmax microplate reader (Molecular Devices, San Jose, CA). For BrdU immunofluorescence staining, SMCs were seeded in 8-chamber slides (MatTek, Ashland, MA), starved, and subsequently stimulated with 10% FBS for 24 h. 40 µM BrdU was added 3 h before the end of the experiment. Cells were fixed with 4% paraformaldehyde (4% PFA), permeabilized with 0.2% Triton X-100 in PBS, and blocked with Dako serum-free blocking solution (Dako, X090930). Immunofluorescence staining was performed using a primary antibody against BrdU (Bio-Rad, MCA2483T) followed by a secondary antibody Alexa Fluor 488 (ThermoFisher Scientific). Nuclei were stained with DAPI. Images were acquired using a Zeiss AxioImager M1 microscope.

### siRNA Transfection

Rat MMP-3 siRNA and scrambled siRNA were synthesized by Millipore Sigma (Burlington, MA). Human MMP-3 siRNA pool and control siRNA pool were purchased from Dharmacon (Lafayette, CO). Rat and human vein SMCs were transfected with siRNA using Lipofectamine RNAi/MAX reagent (Thermo Fisher Scientific, 13778-075) according to the manufacturer’s instructions.

### Electroporation

The plasmid pCMV3-MMP3-Flag was purchased from Sino Biological (Wayne, PA). The constitutively active Akt plasmid pcDNA3.1-Myr-HA-Akt1 and the dominant-negative Akt plasmid HA-AKT-DN (K179M) were obtained from Addgene (Watertown, MA). For electroporation transfection, rat vein SMCs were trypsinized, washed with PBS, and resuspended in cold electroporation buffer. A 250 µl aliquot of SMC suspension (1x10^6^ cells) and 15 µg of plasmid DNA were added to a 4-mm electrode gap cuvette (Bio-Rad, Hercules, CA), mixed thoroughly, and subjected to a single pulse at 300 V and 500 µF using a Gene Pulser electroporator (Bio-Rad, Hercules, CA). Following electroporation, SMCs were transferred to 6-well or 96-well culture plates containing complete DMEM medium and incubated in a humidified incubator (37°C, 5% CO2). The culture medium was replaced with fresh medium the following day.

### Quantitative real-time PCR

Total RNA was isolated from rat and human vein SMCs using RNeasy Mini Kit (Qiagen, Hilden, Germany). Frozen AVF and control veins from mice and human specimens were ground on dry ice, followed by RNA isolation using RNeasy Mini Kit. Complementary DNA (cDNA) was synthesized using the iScript cDNA Synthesis Kit (Bio-Rad, 170-8890). Quantitative polymerase chain reaction (qPCR) amplifications were performed using PowerUp SYBR Green Master Mix (Thermo Fisher Scientific, A25742) in a Bio-Rad CFX96 Real-Time PCR System. Relative mRNA levels were quantified using the comparative Ct method and normalized to the internal control glyceraldehyde-3-phosphate dehydrogenase (GAPDH). The primers used for qPCR are listed in Table 1 of the Supplemental Materials.

### Western blotting

Vein SMCs and tissue powder were lysed in RIPA buffer with a protease inhibitor cocktail (Millipore Sigma, P8340). The concentrations of total proteins were measured using Pierce BCA Protein Assay Kit (ThermoFisher Scientific, 23225). Briefly, 5-20 μg of protein lysates was loaded on 10% SDS-PAGE and electro-transferred into PVDF membrane at 4 °C overnight. The membranes were blocked with 5% milk for 1.5 h, incubated with primary antibodies at room temperature for 2 h, and then horseradish peroxidase-coupled secondary antibodies at room temperature for 1 h. The signals were visualized using Pierce™ ECL Western Blotting Substrate (Thermo Fisher Scientific, 32106). The primary antibodies for Western blotting were listed in Supplemental Table 2.

### Cell cycle analysis

The distribution of vein SMCs in the cell cycle was assessed using flow cytometry. To evaluate the effect of MMP-3 siRNA on cell cycle progression, SMCs were transfected with 100 nM MMP- 3 siRNA or scrambled siRNA. After a 2-day serum-free starvation, the cells were stimulated with 5% FBS for 24 h. To assess the effect of MMP-3 inhibition on cell cycle progression, SMCs underwent serum-free starvation, followed by stimulation with 5% FBS in the presence or absence of the MMP-3 inhibitor for 24 h. At the end of the experiments, cells were fixed with 70% ethanol/PBS at 4 °C, treated with a 10 µg/ml RNase A solution at 37 °C for 15 min, and then stained with 10 µg/ml propidium iodide at room temperature in the dark for 0.5 h. Flow cytometry analysis was conducted using a cytoFLEX flow cytometer (Beckman Coulter Life Sciences, Indianapolis, IN), and the cell population in each phase of the cell cycle was calculated utilizing FlowJo v. 10.8.1 Cell cycle Platform with Watson (pragmatic) model.

### Immunofluorescent staining

Rat vein SMCs were washed with PBS and fixed with 4% paraformaldehyde (PFA). The AVF was dissected, embedded in paraffin, and cut into 5 μm cross-sections. The sections were deparaffinized and treated with Antigen Unmasking Solution (Vector Laboratories, H-3300) in 95 °C water bath for 30 min for antigen retrieval. SMCs or tissue sections were permeabilized in 0.2% Triton x-100/PBS, and then blocked with Dako serum-free blocking solution (Dako, X090930) and incubated with primary antibody at 4 °C overnight, followed by incubation of secondary antibody Alexa Fluor 546 or 488 (Thermo Fisher Scientific). Nuclei were stained with DAPI. The staining was visualized with a Zeiss AxioImager M1 microscope. The primary antibodies for immunofluorescence staining were listed in Supplemental Table 2.

### Statistical analysis

Data are represented as mean ± SD or mean ± SE. Statistical analyses were performed using GraphPad Prism 9.0 software. All in vitro experiments were independently repeated at least three times. Quantitative analyses of microscopic images were conducted in a blinded manner. Sample size calculations for each *in vivo* experiment were performed using PASS software 2019. For comparisons between two unpaired groups, Student’s t-test was used. For comparisons among three or more groups, one-way ANOVA or two-way ANOVA adjusted with Tukey’s post-hoc test was used. The patency of AVF will be analyzed using Kaplan Meier analysis. The Log-rank test will be used for patency curve comparison. The p-value of less than 0.05 was considered statistically significant.

## Results

### MMP-3 is highly induced during AVF neointimal hyperplasia *in vivo*

We used a mouse aortocaval AVF model to determine whether MMP-3 was induced during AVF- associated neointimal development. This is a well-established model that can be used to study both AVF maturation and failure.^38–40, 43, 44^ We have previously shown that AVF begins to fail after a 21-day maturation phase.^45^ Therefore, we collected the inferior vena cava (IVC) and abdominal aorta at day 21 after AVF creation and examined MMP-3 levels using qPCR and Western blotting. As expected, cell proliferation marker PCNA was increased in the IVC but was not changed in the abdominal aortas after AVF creation (**Fig. 1A-D**). Accordingly, MMP-3 mRNA and protein levels were markedly increased in the IVC after AVF. Our previous data showed that neointimal hyperplasia in the outflow vein and the juxta-anastomotic area (JAA) contributes to AVF neointimal failure.^43, 46^ To further examine the MMP-3 expression pattern, we used immunofluorescence double staining for MMP-3 and SMC marker SM-α-actin in the outflow IVC. MMP-3 expression was low in the IVC of the Sham-treated mice, however, it was markedly increased in animals with AVF, which largely colocalized with SM-α-actin in the neointimal area of the IVC (**Fig. 1E-F**). Additionally, we observed that MMP-3 was increased in the JAA after AVF (**Fig. 1G-H**). These data suggest that SMC MMP-3 could be involved in AVF neointimal failure *in vivo*.

**Fig. 1.**
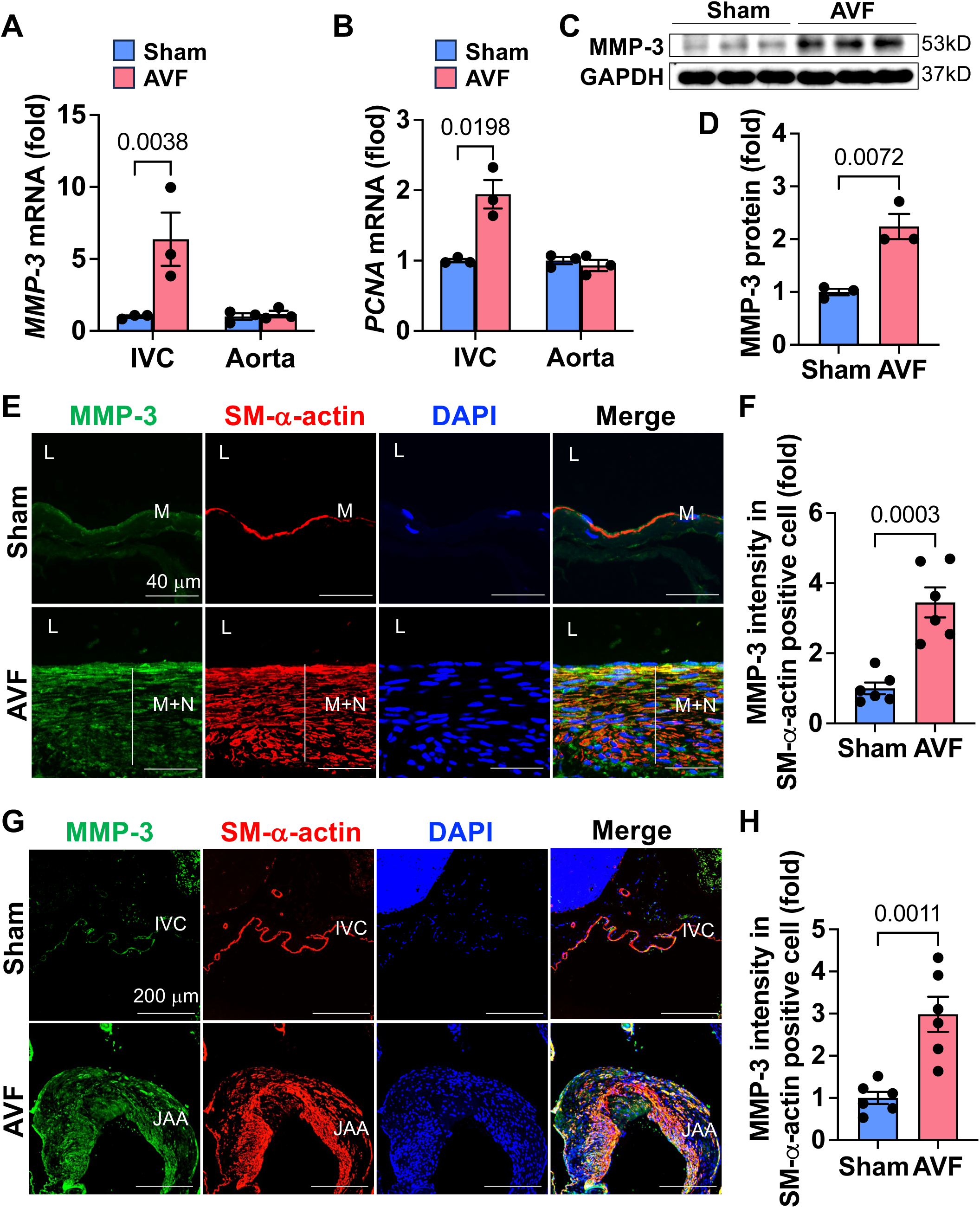
MMP-3 is highly induced during AVF neointimal failure *in vivo*. 9-week-old male C57BL/6J mice underwent an arteriovenous fistula (AVF) creation procedure by puncturing the abdominal aorta and the inferior vena cava (IVC) using a 25-gauge needle. The animals were harvested after 21 days. **A-B**, qPCR data showed that the expression of MMP-3 was exclusively upregulated in the IVC but not in the aortas following AVF creation. As expected, the proliferative gene *PCNA* was also increased in the IVC of these animals after AVF. n=3. **C-D**, Western blotting showed that MMP-3 was upregulated in the IVC after AVF. n=3. **E**, Immunofluorescence double staining showed that MMP-3 was markedly increased in the neointimal area of the outflow IVC in AVF mice compared with the Sham controls, which co-localized with SMC marker SM-α-actin. n=6. **F**, Quantitative results of immunofluorescence staining in the outflow IVC. **G**, Immunofluorescence double staining showed that MMP-3 upregulated in growing SMC in the juxta-anastomotic area (JAA) after AVF. **H**, Quantitative results of immunofluorescence staining in the JAA. n=6. qPCR results were normalized with the internal control gene *GAPDH*. For qPCR analysis, due to tiny tissue, three veins from 3 mice were pooled together as one sample. Values are mean ± SE. Data were analyzed using an unpaired Student’s t-test. *P*<0.05 was significant. **M**, media; **N**, neointima; **L**, lumen; **DAPI**, nucleus; **IVC,** inferior vena cava.

### MMP-3 is highly expressed in AVF neointimal hyperplasia in patients with ESKD

To investigate whether MMP-3 is also increased in human AVF specimens, we collected control and basilic veins with significant neointimal hyperplasia from patients with ESKD during the second stage basilic vein transposition. Immunofluorescence double staining showed that MMP- 3 was highly expressed in the neointimal area (white arrows) of the AVF veins, which largely overlapped with SMC marker SM-α-actin in the neointimal area (**Fig. 2A-B**). Immunofluorescence staining for cell proliferation marker PCNA further showed that MMP-3 was highly expressed in the growing SMCs (**Fig. 2C-D**). qPCR and Western blotting results showed that both MMP-3 and PCNA were significantly increased in the AVF compared with the control veins (**Fig. 2E-H**). These data suggest that MMP-3 was primarily increased in the neointimal SMC and may play a role in human AVF neointimal hyperplasia.

**Fig. 2.**
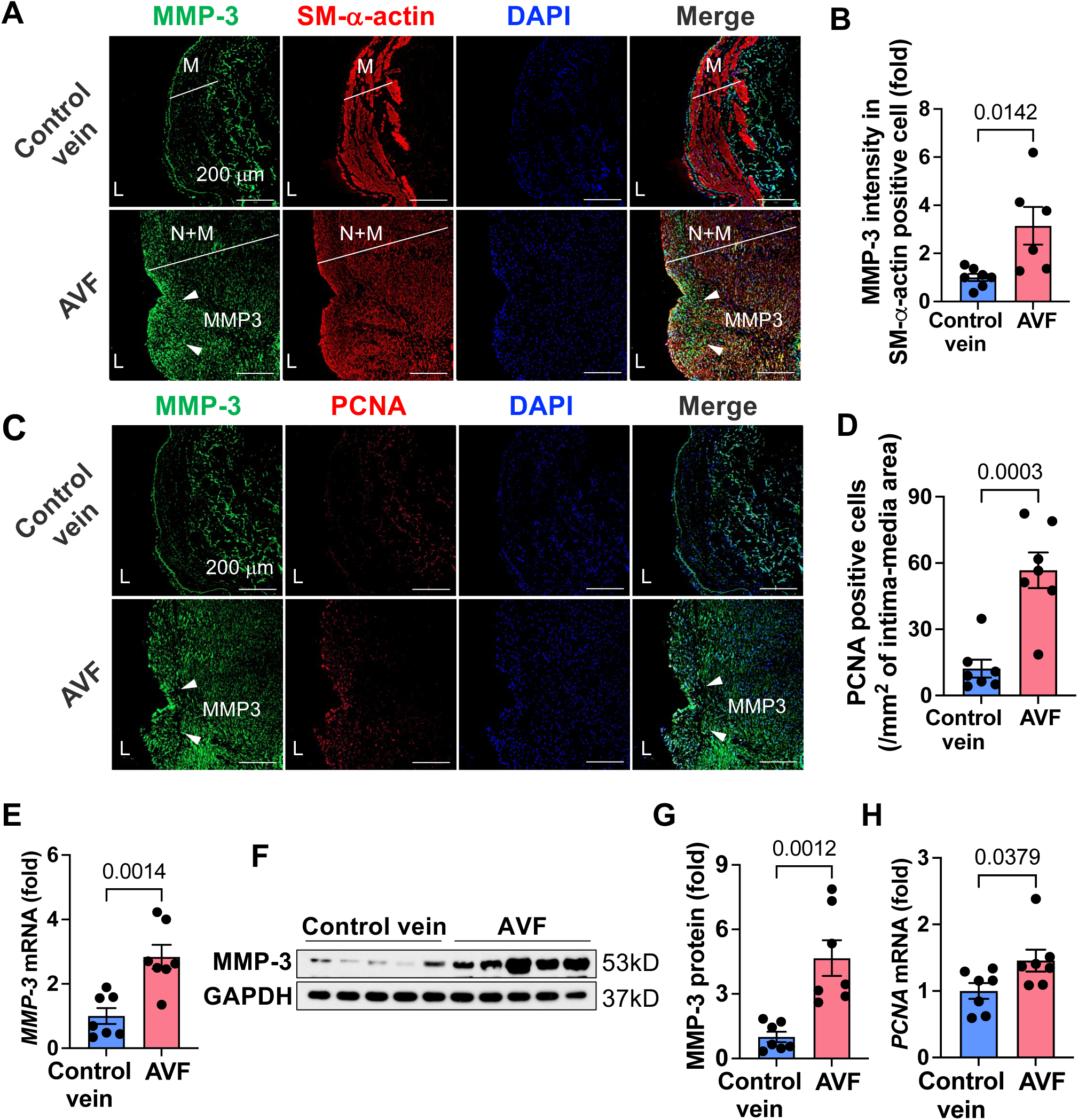
MMP-3 is highly expressed in human AVF specimens from patients with ESKD. Human AVF specimens and control veins were from patients with ESKD undergoing a second stage of basilic vein transposition procedure. **A-D,** Immunofluorescence double staining showed that MMP-3 was highly expressed in the AVF veins and largely overlapped with SMC markers SM-α-actin and proliferative marker PCNA in the neointimal area. **B** and **D**, Quantitative data of immunofluorescence staining for SM-α-actin and PCNA. n=6-7. **E-G,** qPCR and Western blotting showed that MMP-3 expression was higher in AVF veins than the control veins. n=7. **H**, The proliferative gene *PCNA* was increased in the AVF veins compared with the controls. n=7. qPCR results were normalized with the internal control gene *GAPDH*. Values are mean ± SE. Data were analyzed using unpaired Student’s t-test. *P*<0.05 was significant. **M**, media; **N**, neointima; **L**, lumen; **DAPI**, nucleus; White arrows, MMP-3.

### Knockdown and inhibition of MMP-3 suppress SMC proliferation and ECM deposition *in vitro*

Since SMC proliferation significantly contributes to AVF neointimal failure, we determined whether MMP-3 plays a role in SMC proliferation. We first examined whether MMP-3 was induced in response to FBS and the growth factor PDGF-BB. Both FBS and PDGF-BB time-dependently upregulated MMP-3, and the peak time of MMP-3 induction was 3 h (**Fig. 3A**). To determine whether MMP-3 is required for venous SMC proliferation, the specific MMP-3 siRNA was used. As expected, FBS markedly increased cell proliferation in scrambled siRNA-treated rat vein SMC, however, this effect was significantly reduced in MMP-3 siRNA-treated SMC (**Fig. 3B-D**). Furthermore, we observed that MMP-3 inhibitor dose-dependently suppressed FBS-stimulated SMC proliferation (**Fig. 3E**). To examine whether MMP-3 mediates venous SMC proliferation due to specifically targeting MMP-3, we overexpressed MMP-3 in rat venous SMC (**Fig. 3F**), and MMP-3 overexpression significantly promoted SMC proliferation (**Fig. 3G**). Next, we examined whether MMP-3 was required for human venous SMC proliferation. Both FBS and PDGF-BB also time-dependently increased MMP-3 expression in human vein SMC (**Fig. 3H**). Likewise, MMP-3 siRNA and inhibitor significantly attenuated FBS-stimulated SMC proliferation (**Fig. 3I-L**). Deposition of extracellular matrix (ECM) plays an important role in mediating SMC proliferation and neointimal hyperplasia,^2, 47^ and fibronectin and collagen are the most common ECM proteins involved in pathological vascular remodeling. Therefore, we examined the role of MMP-3 in ECM deposition. FBS markedly increased intracellular and extracellular collagen and fibronectin in scrambled siRNA-treated SMC; however, these effects were antagonized in MMP-3 siRNA- treated cells (**Fig. 3M-R**). These data suggest that MMP-3 plays a key role in mediating venous SMC proliferation and ECM deposition.

**Fig. 3.**
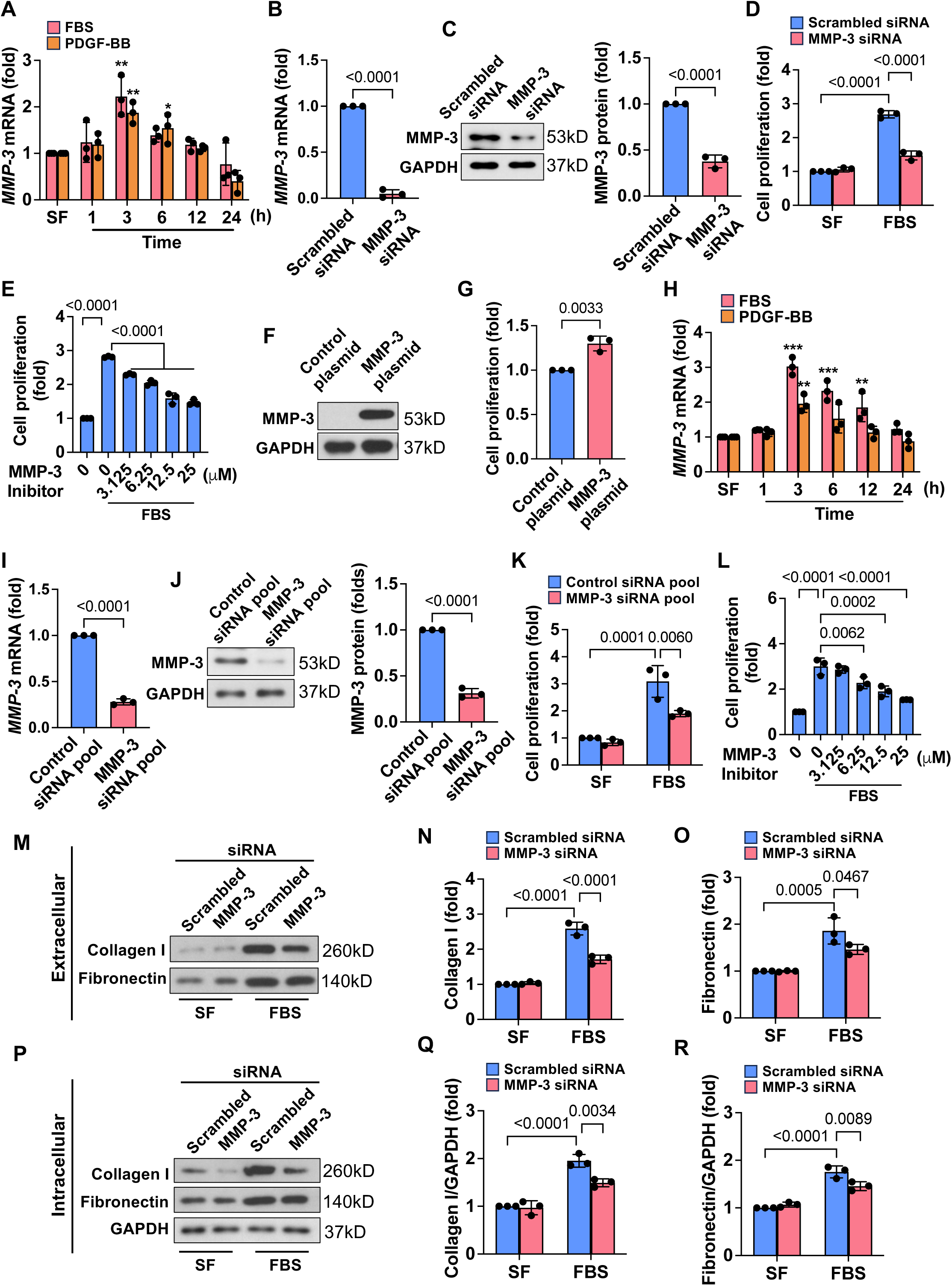
Knockdown and inhibition of MMP-3 suppress SMC proliferation and ECM deposition *in vitro.* **A**, qPCR results showed that MMP-3 was time-dependently induced by FBS and PDGF-BB in venous SMCs. Rat vein SMCs were serum-free starved, stimulated with 10% FBS or 20 ng/ml PDGF-BB, and then harvested at the different time points. n=3. **B-C**, MMP-3 siRNA largely decreased MMP-3 mRNA and protein levels. Rat vein SMCs were transfected with 100 nM scrambled siRNA or MMP-3 siRNA for 48 h. n=3. **D**, Knockdown of MMP-3 inhibited FBS- stimulated SMC proliferation. Rat vein SMCs were transfected with 100 nM scrambled siRNA or MMP-3 siRNA, serum-free starved, followed by stimulation with 10% FBS for 48 h. n=3. Cell proliferation was examined using SRB assay. **E**, SRB assay showed that MMP-3 inhibitor dose- dependently suppressed FBS-induced SMC proliferation. Rat vein SMCs were serum-free starved and pre-treated with indicated concentrations of MMP-3 for 0.5 h, followed by stimulation with 10% FBS for 48 h. n=3. **F**, Western blotting showed that MMP-3 overexpression increased the expression of MMP-3. Rat vein SMCs were transfected with 15 mg of pCMV3-MMP3-Flag or control plasmid using electroporation for 3 days. **G**, SRB assay showed that overexpression of MMP-3 promoted SMC proliferation. Rat vein SMCs were transfected with 15 ug pCMV3-MMP3- Flag or control plasmid using electroporation and then cultured in a DMEM medium containing 0.5% FBS for 48 h. n=3. **H**, qPCR data showed that FBS and PDGF-BB time-dependently increased MMP-3 expression in human venous SMCs. Human vein SMCs were serum-free starved and then stimulated with 10% FBS or 20 ng/ml PDGF-BB for indicated time points. n=3. **I-J**, Western blotting showed that MMP-3 siRNA pool decreased MMP-3 expression in human vein SMCs. Human vein SMCs were transfected with 100 nM scrambled siRNA pool or MMP-3 siRNA pool for 48 h. n=3. **K**, SRB assay showed that knockdown of MMP-3 markedly suppressed SMC proliferation in human vein SMCs. **L**, SRB assay showed that MMP-3 inhibitor dose- dependently attenuated FBS-stimulated SMC proliferation in human vein SMCs. n=3. **M-O**, Effect of MMP-3 siRNA on FBS-induced extracellular collagen I and fibronectin. n=3. **P-R**, Effect of MMP-3 siRNA on FBS-induced intracellular collagen I and fibronectin. n=3. Rat vein SMCs were transfected with 100 nM scrambled siRNA or MMP-3 siRNA, and serum-free starved, followed by stimulation with 5% FBS for 24 h. Both SMCs and culture media were collected, and Western blotting was performed. Values are mean ± SE. Data were analyzed using Student’s t-test, one-way or two-way ANOVA adjusted with Tukey’s post-hoc test for multiple comparisons. *P*<0.05 was significant.

### Global MMP-3 deficiency decreases neointimal hyperplasia and improves AVF patency

To determine the role of MMP-3 in AVF neointimal hyperplasia and failure, MMP-3-WT and MMP- 3-KO mice underwent AVF creation and were harvested after 21 or 42 days (**Fig. 4A**). There was no wall thickening or neointimal hyperplasia in the outflow IVC of MMP-3-WT and MMP-3-KO mice in the Sham group; however, after AVF creation, the wall thickness and neointima + media area were robustly increased in MMP-3-WT mice but these were diminished in MMP-3-KO mice at both days 21 and 42 (**Fig. 4B-F**). Data from Doppler ultrasound showed that the IVC diameter of MMP-3-WT and MMP-3-KO mice was small in the Sham group but increased in response to AVF creation; no significant difference in the diameter was observed between MMP-3-WT and MMP-3-KO mice after AVF (**Suppl. Fig. 1A-B**). Hemodynamic analysis showed that blood flow, velocity, and shear stress were expectedly increased in the IVC of MMP-3-WT and MMP-3-KO mice after AVF; however, there were no significant changes observed between these two groups. Next, we examined the effect of MMP-3 deficiency on neointimal hyperplasia in the JAA. Deletion of MMP-3 markedly reduced neointimal area in the JAA at both days 21 and 42 after AVF (**Fig. 4G-I**). More importantly, ultrasound showed that deletion of MMP-3 significantly improved AVF patency (**Fig. 4J**). Additionally, we determined whether MMP-3 regulated AVF neointimal SMC proliferation *in vivo*. Immunofluorescence double staining showed that the proliferation marker Ki67 was increased in the neointimal SMCs in the outflow IVC in MMP-3-WT mice after AVF; however, these effects were blunted in MMP-3-KO mice (**Fig. 4K-L**). Similarly, we observed that deficiency of MMP-3 decreased SMC proliferation in the JAA after AVF (**Fig. 4M-N**). These data suggest that MMP-3 plays a causative role in AVF neointimal failure.

**Fig. 4.**
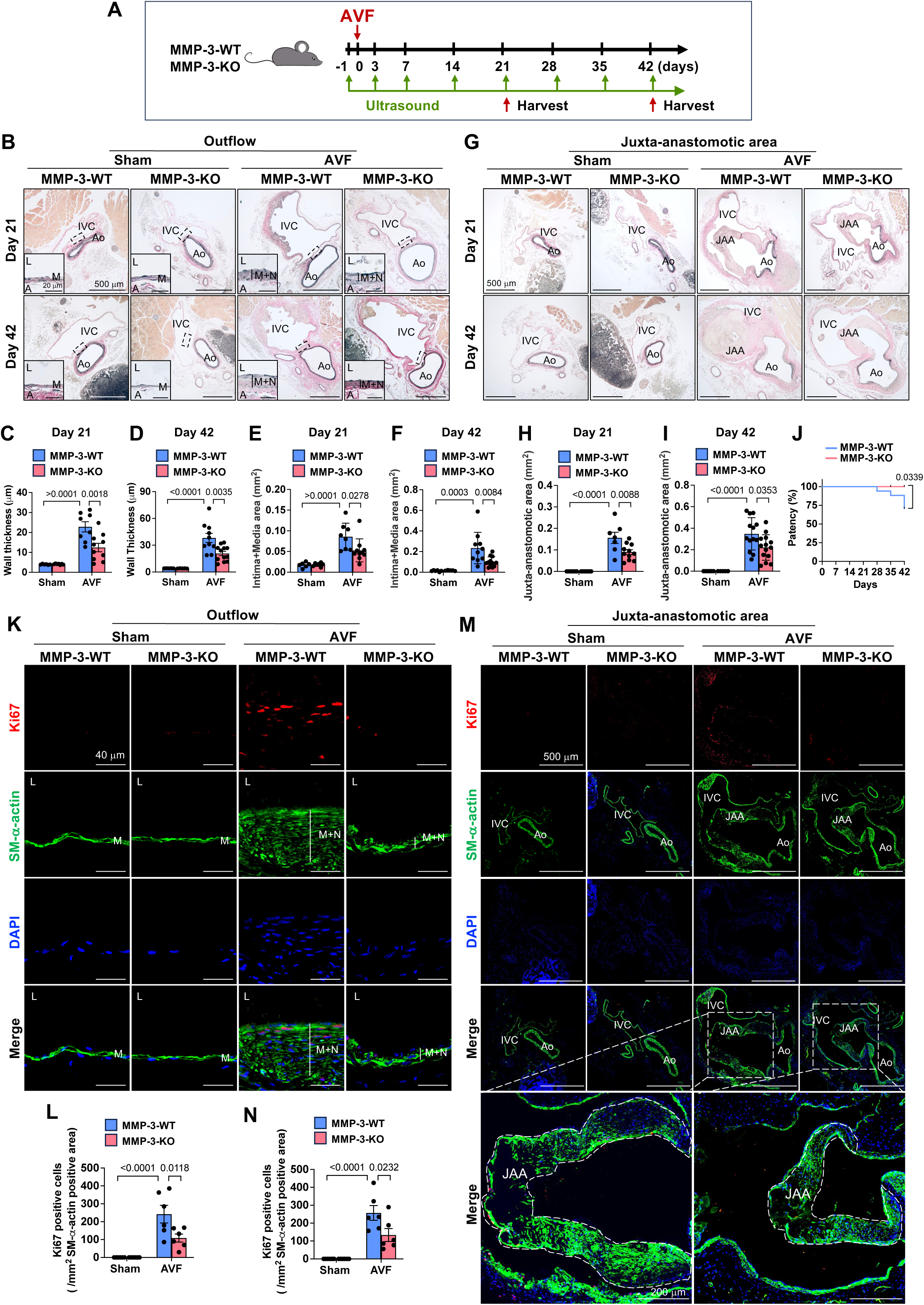
Global MMP-3 deficiency decreases neointimal hyperplasia and improves AVF patency. **A**, Experimental design. 9-week-old male MMP-3-WT and MMP-3-KO mice were subjected to AVF creation or Sham surgery and harvested after 21 days and 42 days. **B**, MMP-3- KO mice had less wall thickness and neointima + media area in the outflow IVC than MMP-3-WT mice after AVF. n=5-14. The elastic lamellae were determined using VVG staining. **C-D**, Quantitative data of wall thickness. **E-F**, Quantitative data of intima + media area. **G**, VVG staining showed that MMP-3-KO mice had less juxta-anastomotic area (JAA) of the IVC than MMP-3-WT mice after AVF. n=6-12. **H-I**, Quantitative data of the JAA. **J**, Deficiency of MMP-3 improved IVC patency after AVF. The patency was monitored by ultrasound. n=14-17. **K**, Representative immunofluorescence images showed that MMP-3-KO mice had lower proliferation marker PCNA than MMP-3-WT mice in the neointima of the outflow IVC after AVF. **L**, Quantitative results of PCNA and SM-α-actin in the outflow IVC. n=6. **M**, Representative immunofluorescence images showed that MMP-3-KO mice had lower proliferation marker PCNA than MMP-3-WT mice in the JAA of the IVC after AVF. **N**, Quantitative results of PCNA and SM-α-actin in the JAA of the IVC. n=6. Values are mean ± SE. Data were analyzed using two-way ANOVA adjusted with Tukey’s post-hoc test for multiple comparisons. Kaplan-Meier (Log rank) test was used for AVF patency analysis. *P*<0.05 was significant. The selected area was enlarged for a better view. **IVC,** inferior vena cava; **Ao**, aorta; JAA, juxta-anastomotic area; **M**, media; **N**, neointima; **L**, lumen; **DAPI**, nucleus.

### Deficiency of SMC-specific MMP-3 reduces neointimal hyperplasia and improves AVF patency

Since MMP-3 was increased in the neointimal SMCs after AVF and knockdown of MMP-3 was able to inhibit SMC proliferation *in vitro*, we next investigated whether MMP-3 deletion in SMC is sufficient to reduce AVF neointimal hyperplasia and failure using SMC-specific MMP-3-KO mice. SMC-specific MMP-3-KO mice were generated as we previously described.^23^ We further validated these mice in the AVF for 21 days using immunofluorescence staining. In Oil-injected SMMHC-Cre^+^/MMP-3-flox^+/+^ mice with AVF, MMP-3 was expressed in the neointimal area of the IVC, as well as in the aorta, which colocalized with SM-α-actin; however, after TMX injection, MMP-3 was undetectable in these areas, indicating that MMP-3 was genetically deleted in SMCs in these mice (**Suppl. Fig. 2**). We next determined whether MMP-3 in SMC was required for AVF failure. SMMHC-Cre^+^/MMP-3-flox^+/+^ mice were intraperitoneally (IP) injected with Oil or TMX and then underwent AVF creation and harvested after 42 days (**Fig. 5A**). There was no difference in the wall thickness and intima + media area in the outflow IVC between Oil- and TMX-injected Sham mice; in contrast, AVF creation resulted in significant wall thickening and neointimal hyperplasia in Oil-injected mice, and these effects were significantly reduced in TMX-injected mice (**Fig. 5B - D**). Moreover, we observed that the JAA was markedly increased in Oil-injected mice after AVF but was significantly decreased in TMX-injected mice (**Fig. 5B** and **E**). Additionally, we observed that SMC-specific MMP-3 deletion also significantly reduced the proliferation marker Ki67 in the neointimal SMCs in both the outflow and JAA of the IVC of TMX-injected SMMHC-Cre^+^/MMP-3- flox^+/+^ mice after AVF (**Fig. 5G-J**). These data suggest that MMP-3 in SMCs is a key mediator of AVF neointimal failure.

**Fig. 5.**
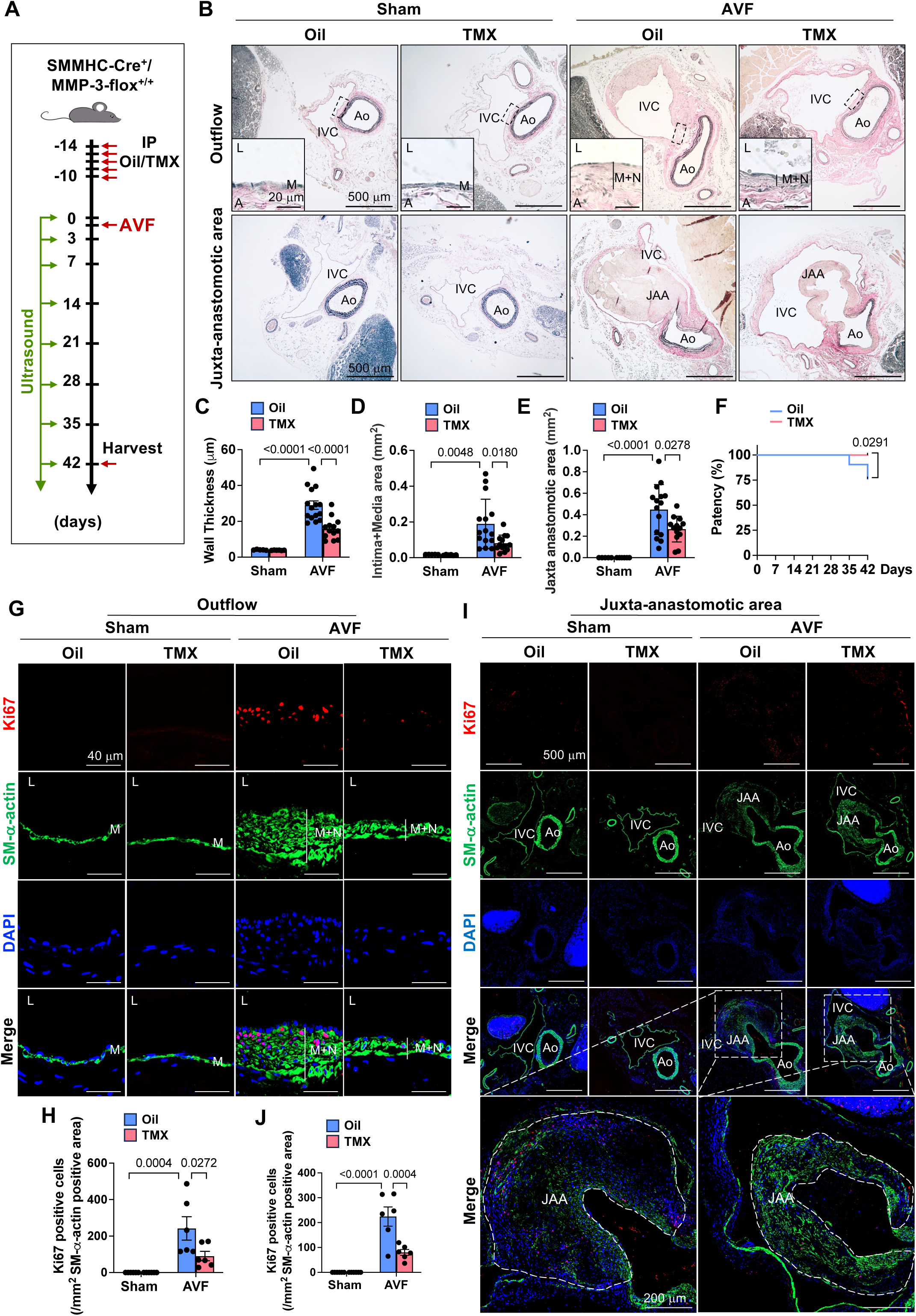
Deficiency of SMC-specific MMP-3 reduces neointimal hyperplasia and improves AVF patency. **A**, Experimental design. 7-week-old male SMMHC-Cre^+^/MMP-3-flox^+/+^ mice were IP injected with 75 mg/kg tamoxifen (TMX) or Oil for 5 consecutive days, followed by 2 weeks of rest, and then subjected to AVF creation. Mice were harvested after 42 days. **B**, SMC-specific MMP-3 deficiency (TMX injection) reduced wall thickness and neointima + media area in the outflow IVC and the JAA compared with wild-type (Oil injection) after AVF. n=5-15. **C**, Quantitative results of wall thickness. **D**, Quantitative results of intima + media area. **E**, Quantitative data of the JAA. **F**, Deletion of MMP-3 in SMCs improved the patency of the IVC after AVF. Ultrasound was used to monitor AVF patency. n=18-21. **G-H**, Immunofluorescence staining showed that SMC-specific MMP-3 deficiency decreased the proliferation marker PCNA in the outflow IVC compared with wild-type after AVF. n=6. **I-J**, Effect of SMC-specific MMP-3 deficiency on PCNA in the JAA. n=6. Values are mean ± SE. Data were analyzed using two-way ANOVA adjusted with Tukey’s post-hoc test for multiple comparisons. Kaplan-Meier (Log rank) test was used to analyze AVF patency. *P*<0.05 was significant. The selected area was enlarged for a better view. **IVC,** inferior vena cava; **Ao**, aorta; JAA, juxta-anastomotic area; **M**, media; **N**, neointima; **L**, lumen; **DAPI**, nucleus.

### Knockdown and inhibition of MMP-3 arrest venous SMC at the G1 phase

Since knockdown and inhibition of MMP-3 suppress SMC proliferation, we next examined the role of MMP-3 in cell cycle progression. As shown in **Fig. 6A-B**, the data from flow cytometry showed that approximately, 80% of cells were in the G1 phase when rat vein SMC were serum-free starved. In response to FBS stimulation, the percentage of cells in the G1 phase was markedly decreased (about 57%), whereas the percentage of the cells in the S and G2 phases was increased. However, MMP-3 siRNA significantly reduced cell percentage in the S and G2 phases and increased the cells in the G1 phase compared with control siRNA after FBS stimulation. Moreover, MMP-3 inhibitor also dose-dependently blocked FBS-induced cell accumulation in the S and G2 phases and arrested the cells in the G1 phase. (**Fig. 6C**). These data suggest that knockdown and inhibition of MMP-3 suppress SMC proliferation via cell cycle G1 arrest. Cell cycle progression is driven by the cyclin/CDK (cyclin-dependent kinase) complexes,^48, 49^ and p21^CIP1^/p27^KIP1^ are CDK inhibitors that can bind to the cyclin/CDK complexes to induce cell cycle arrest.^50^ Therefore, we examined the expression of the cyclins, p21^CIP1^, p27^KIP1^, and cell proliferation marker PCNA using qPCR. As expected, MMP-3 siRNA suppressed FBS-induced SMC proliferation showing decreased PCNA expression (**Fig. 6D**). Accordingly, knockdown of MMP-3 suppressed FBS-induced upregulation of the cyclin B1, D1, and E1 (**Fig. 6E-G**). Additionally, we showed that MMP-3 siRNA blocked FBS-induced downregulation of p21^CIP1^ and p27^KIP1^ (**Fig. 6H-I**). These data suggest that MMP-3 mediates SMC proliferation at least in part via regulating p21^CIP1^ and p27^KIP1^.

**Fig. 6.**
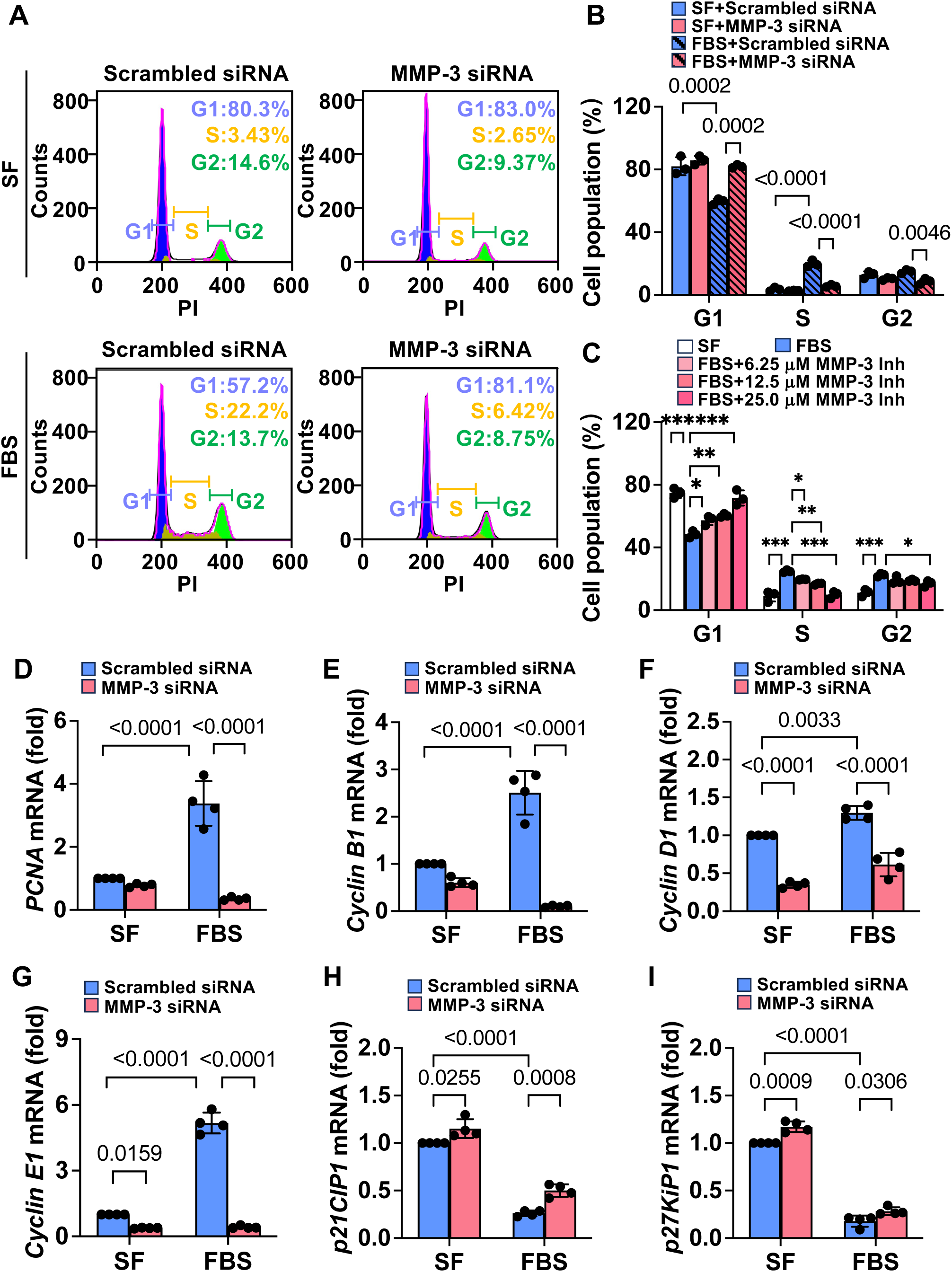
Knockdown and inhibition of MMP-3 arrest venous SMC at the G1 phase. **A,** Representative flow cytometry analysis showing the effects of MMP-3 siRNA on cell cycle progression. Rat vein SMCs were transfected with 100 nM scrambled siRNA or MMP-3 siRNA, and then serum-free starved, followed by stimulation with 5% FBS for 24 h. **B**, Quantitative results of flow cytometry analysis. n=3. **C**, MMP-3 inhibitor dose-dependently blocked FBS-induced cell cycle progression. Rat vein SMCs were serum-free starved, and then treated with indicated concentrations of MMP-3 inhibitor, followed by stimulated with 5% FBS for 24 h. n=3. MMP-3 inh, MMP-3 inhibitor. **D–I**. qPCR data showing the effects of MMP-3 knockdown on PCNA, cyclin B1, D, E1, p21^CIP1^, and p27^KIP1^ during SMC proliferation. Rat vein SMCs were transfected with 100 nM scrambled siRNA or MMP-3 siRNA, and then serum-free starved, followed by stimulated with 5% FBS for 24 h. qPCR results were normalized with the internal control gene *GAPDH*. n=4. Values are mean ± SD. Data were analyzed using one-way or two-way ANOVA adjusted with Tukey’s post-hoc test for multiple comparisons. \**P* < 0.05, \*\**P* < 0.01, \*\*\**P* < 0.001, compared with FBS. *P*<0.05 was significant.

### MMP-3 mediates venous SMC proliferation via regulating FAK-AKT signaling

The kinases FAK, AKT, and ERK play critical roles in SMC proliferation and neointimal hyperplasia;^33, 42, 51, 52^ therefore, we determined whether MMP-3 regulates these kinases in venous SMC. FBS stimulation increased p-FAK, p-AKT, and p-ERK; however, MMP-3 siRNA significantly attenuated FBS-induced p-FAK and p-AKT but did not affect p-ERK (**Fig. 7A-D**). Overexpression of MMP-3 significantly increased p-FAK and p-AKT (**Fig. 7E-G**), suggesting that MMP-3 activates FAK and AKT during SMC proliferation. Since nuclear FAK can inhibit SMC proliferation, and therefore suppress neointimal hyperplasia,^33, 34^ we next determined whether MMP-3 regulates FAK nuclear-cytoplasm translocation in rat vein SMC. Immunofluorescence staining showed that knockdown of MMP-3 markedly mediated FAK nuclear translocation (**Fig. 7H-I**). To determine whether FAK signaling is critical for MMP-3-mediated SMC proliferation, we used a FAK inhibitor VS-4718;^33, 34^ treatment with VS-4718 dose-dependently inhibited FBS-induced SMC growth, suggesting the importance of FAK in SMC proliferation (**Fig. 7J**). Furthermore, VS-4817 treatment blocked MMP-3 overexpression-induced SMC proliferation (**Fig. 7K**), suggesting that FAK signaling is required for MMP-3-mediated SMC proliferation. To determine whether activation of AKT is essential for MMP-3-induced SMC proliferation, we used constitutively activated AKT (CA- AKT) and dominant-negative AKT (DN-AKT) constructs. As shown in **Fig. 7L-M**, transfection of CA-AKT largely increased both p-AKT and total AKT in rat vein SMC, indicating that AKT was activated. Accordingly, we found that CA-AKT transfection significantly blocked MMP-3 siRNA- inhibited SMC proliferation. Conversely, DN-AKT transfection suppressed MMP-3 overexpression-induced SMC proliferation (**Fig. 7N-O**). These data suggest that AKT activation is required for MMP-3-mediated SMC proliferation. Emerging evidence has shown that FAK could be an upstream of AKT.^36, 37, 53^ Therefore, we further determined whether MMP-3 mediated SMC proliferation via FAK-AKT signaling. As shown in **Fig. 7P-R**, as expected, FAK inhibitor VS-4718 not only suppressed MMP-3 overexpression-induced p-FAK but also attenuated p-AKT, suggesting that AKT could be a downstream regulator of FAK in MMP-3-mediated SMC proliferation. Additionally, we examined p-FAK and p-AKT in MMP-3-WT and MMP-3-KO mice after AVF *in vivo*. As shown in **Fig. 7S-V**, immunofluorescence staining showed that the intensity of p-FAK and p-AKT was increased in the neointimal SMC of the outflow IVC of MMP-3-WT mice after AVF; however, these effects were significantly blocked in MMP-3-KO mice. These data suggest that FAK-AKT signaling plays a critical regulatory role in MMP-3-mediated SMC proliferation.

**Fig. 7.**
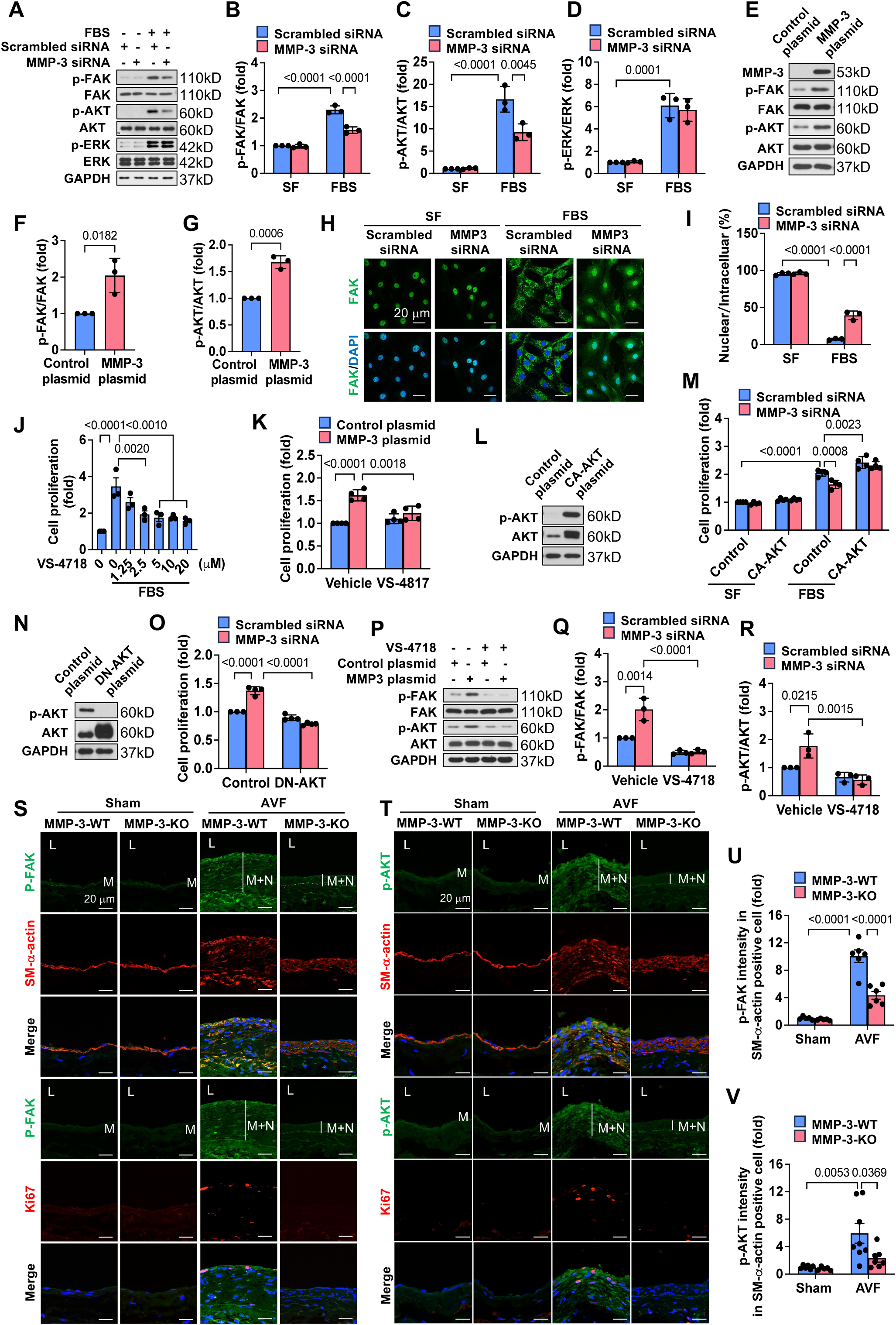
MMP-3 mediates venous SMC proliferation via regulating FAK-AKT signaling. **A**, Western blotting showing the effects of MMP-3 siRNA on FBS-induced p-FAK, p-AKT and p-ERK. Rat vein SMCs were transfected with 100 nM scrambled siRNA or MMP-3 siRNA, and then serum-free starved, followed by stimulated with 10% FBS for 24 h. n=3. **B-D**, Quantitative results of p-FAK, p-AKT and p-ERK. Data were normalized with total FAK, AKT or ERK. **E**, Western blotting showed that the overexpression of MMP-3 increased p-FAK and p-AKT expression. Rat vein SMCs were transfected with pCMV3-MMP3-Flag or control plasmid using electroporation and then cultured in 0.5% FBS DMEM medium for 48 h. n=3. **F-G**, Quantitative results of p-FAK and p-AKT. **H**, Immunofluorescence staining showed that MMP-3 knockdown resulted in FAK nuclear translocation. Rat vein SMCs were transfected with 100 nM scrambled siRNA or MMP-3 siRNA, and then serum-free starved, followed by stimulation with 10% FBS for 24 h. **I**, Quantitative results of FAK. n=3. **J**, FAK inhibitor VS-4718 dose-dependently attenuated FBS-stimulated SMC proliferation in Rat vein SMCs. Cell proliferation was examined using SRB assay. n=3. **K**, SRB assay showed that FAK inhibitor VS-4718 blocked MMP-3 overexpression-induced SMC proliferation. Rat vein SMCs were transfected with pCMV3-MMP3-Flag plasmid or control plasmid in the presence or absence of 2.5 μM VS-4718 for 48 h. n=4. **L-M**, Effects of constitutively activated AKT on MMP-3 knockdown-suppressed SMC proliferation. Rat vein SMCs were transfected with CA-AKT plasmids or MMP-3 siRNA using electroporation, and then serum-free starved, followed by stimulation with 10% FBS for 48 h. **N-O**, Effects of dominant-negative AKT on MMP-3 overexpression-induced SMC proliferation. Rat vein SMCs were transfected with DN-AKT plasmids or MMP-3 plasmid using electroporation, and then cultured in 0.5% FBS DMEM medium for 48 h. n=4. **P**, Western blotting showing the effects of FAK inhibitor VS-4718 on MMP- 3 overexpression-induced p-FAK and p-AKT. Rat vein SMCs were transfected with MMP-3 and control plasmid, and then treated with 2.5 μM VS-4718, cultured in 0.5% FBS DMEM medium for 48 h. n=3. **Q-R**, Quantitative results of p-FAK and p-AKT. Data were normalized with total FAK or AKT. **S-T**, Immunofluorescence staining showed that p-FAK and p-AKT were increased in the neointimal area in the outflow IVC in MMP-3-WT mice after AVF, which was reduced in MMP-3- KO mice. 9-week-old male MMP-3-WT and MMP-3-KO mice were subjected to AVF creation and harvested after 21 days. **U-V**, Quantitative results of immunofluorescence staining for p-FAK and p-AKT. n=5-7. Values are mean ± SD or mean ± SE. Data were analyzed using Student’s t-test, one-way or two-way ANOVA adjusted with Tukey’s post-hoc test for multiple comparisons. *P*<0.05 was significant. **M**, media; **N**, neointima; **L**, lumen; **DAPI**, nucleus.

## Discussion

Neointimal hyperplasia is the leading cause of AVF failure in patients with ESKD, with stenoses in both the outflow vein and the JAA leading to AVF failure.^3, 43^ Currently, no drug target has been successfully developed to prevent or to treat AVF failure. In this study, we investigate the role of MMP-3 in SMC proliferation, AVF neointimal development, and failure. We observed that MMP-3 was increased in proliferating venous SMCs, in neointimal areas in an AVF mouse model, as well as in human AVF specimens with significant neointimal hyperplasia. Knockdown and inhibition of MMP-3 suppressed SMC proliferation, while MMP-3 overexpression promoted cell proliferation *in vitro*. Global and SMC-specific MMP-3 deficiency reduced neointimal hyperplasia in both the outflow vein and the JAA and improved AVF patency *in vivo*. Mechanistic studies showed that MMP-3 mediates SMC proliferation and AVF neointimal hyperplasia via regulating FAK-AKT signaling.

Increasing evidence has demonstrated that the degrading enzyme MMP-3 is not only implicated in ECM remodeling but also in regulating intracellular signaling pathways during the pathogenesis of vascular disorders.^18, 21, 22^ MMP-3 expression is up-regulated in response to various stimuli including growth and pro-inflammatory cytokines.^54, 55^ Silencing MMP-3 inhibits SMC migration, proliferation, and neointima formation following arterial injury.^56^ Moreover, MMP-3 KO mice have decreased SMC migration, proliferation, and neointimal formation after carotid artery ligation,^57^ suggesting the importance of MMP-3 in arterial pathological remodeling. Interestingly, a previous study showed that overexpression of MMP-3 could suppress human aortic SMC migration and reduce vein graft intimal hyperplasia in a rabbit model.^58^ Unfortunately, this study only used MMP- 3 adenovirus to determine the effect of MMP-3 overexpression in human aortic SMC migration *in vitro* and in a rabbit vein graft disease model *in vivo*, which need to be further validated using other approaches such as knockdown and transgenic mice. We recently observed that MMP-3 was robustly increased (> 180 folds) compared with other MMPs in response to AVF creation in a mouse aortocaval AVF model,^24^ suggesting that MMP-3 is associated with AVF remodeling. In the present study, we showed that knockdown and inhibition markedly suppressed venous SMC proliferation and SMC-specific MMP-3 deficiency significantly reduced neointimal hyperplasia and improved AVF patency, strongly suggesting that SMC-specific MMP-3 is a key mediator of AVF neointimal failure.

FAK signaling plays important roles in cell migration, proliferation, and tissue remodeling. ^25, 59, 60^ FAK can be activated through phosphorylation at tyrosine 397, which contributes to arterial injury- mediated SMC proliferation and ECM deposition.^25, 61–63^ Recent studies have also shown that FAK can traffic between the nucleus and cytoplasm in SMCs; in healthy arteries, FAK is predominantly localized to the nucleus of SMCs and displays low activity, however, FAK can translocate from the nucleus to the cytoplasm and become active after arterial injury.^33^ Inhibition of FAK can cause FAK to accumulate in the nucleus, suppressing SMC proliferation and neointima formation.^35^ In this study, we showed that knockdown of MMP-3 suppressed p-FAK in venous SMCs. We also observed that p-FAK is expressed in the cytoplasm under normal culture conditions, but knockdown of MMP-3 resulted in a translocation of FAK from the cytoplasm to the nucleus, suggesting that MMP-3 is capable of activating FAK in venous SMC. Furthermore, inhibition of FAK using VS-4178 blocked MMP-3 overexpression-mediated SMC proliferation, suggesting that FAK is crucial for MMP-3-mediated venous SMC proliferation. It is known that AKT and ERK1/2 signaling play critical roles in SMC migration and proliferation.^64–68^ Differential effects of targeting AKT and ERK1/2 signaling have been reported. For example, we have shown that vinpocetine, a phosphodiesterase 1 inhibitor, suppresses PDGF-BB-induced phosphorylation of ERK1/2 but not AKT in rat aortic SMCs.^52^ Inhibition of SIK3 using HG-9-91-01 was able to block PDGF-induced activation of AKT but not ERK1/2 in rat aortic SMCs.^42^ In this study, using cultured rat vein SMC, we showed that knockdown of MMP-3 blocked FBS-induced phosphorylation of AKT but not ERK1/2, suggesting that MMP-3 can activate AKT. We additionally showed that DN-AKT suppressed MMP-3 overexpression-induced SMC proliferation, while CA-AKT blocked MMP-3 knockdown-inhibited SMC proliferation, suggesting that AKT activation is required for MMP-3- mediated venous SMC proliferation. AKT is downstream of FAK.^36, 37, 69^ Accordingly, we showed that FAK inhibitor VS-4178 blocked MMP-3 overexpression-induced p-AKT. Collectively, these data suggest that FAK-AKT signaling is critical for MMP-3-regulated SMC proliferation. Of note, there is no clear evidence showing that MMP-3 directly regulates FAK activity. MMPs break down ECM structural proteins like elastin and collagen in the vascular wall after arterial injury, which allows SMC to migrate and proliferate.^14, 16, 18^ ECM such as collagen and fibronectin synthesis could also be initiated by FAK activation.^16, 70^ Additionally, the newly synthesized ECM could promote FAK activation. It has been reported that collagen can activate FAK signaling, which plays a key role in liver fibrosis.^71^ Therefore, MMP-3-triggered ECM remodeling could drive FAK activation. Indeed, we observed that knockdown of MMP-3 markedly decreased ECM collagen and fibronectin deposition in rat vein SMC. We have previously shown that p-AKT is increased after AVF and deficiency of AKT1 significantly reduced AVF remodeling. In this study, we showed that MMP-3 deletion blocked AVF-induced AKT activation.^72^ All these data support the construct that FAK-AKT signaling plays a key role in MMP-3-mediated SMC proliferation and AVF neointimal hyperplasia.

Clinical and laboratory evidence including ours has shown that neointimal hyperplasia in the outflow vein and the JAA primarily contributes to AVF neointimal failure.^3, 4, 43, 46^ We have recently comprehensively studied these two areas of neointimal hyperplasia using two commonly used AVF animal models (rat carotid artery-to-jugular vein anastomosis and mouse aortocaval puncture models); although neointimal hyperplasia in the outflow vein and the JAA participated in AVF failure, the JAA more likely played a predominant role.^3, 43, 73^ Several types of cells including endothelial cells and SMCs are involved during AVF venous remodeling.^74–76^ Endothelial cells are believed to be more involved in the early stage of AVF adaptive remodeling such as AVF maturation. We have shown in our recent study that endothelial cells were lost as early as 6 h in the outflow IVC due to disturbed flow response *in vivo*.^43, 46^ By contrast, SMC may play significant roles in AVF neointimal hyperplasia failure. We have observed that SM-α-actin-positive cells increase and ECM deposits robustly in the outflow IVC and the JAA after 21 days (beginning the failure process) following AVF, supporting that overgrowing SMC and ECM deposition largely contribute to aggressive neointimal hyperplasia. The mouse aortocaval AVF model can be used to study both AVF maturation and failure.^40, 43^ In the clinical setting, blood flow and diameter of AVF are two key factors used to examine AVF maturation.^77^ In the present study, we found that both global and SMC-specific MMP-3 deficiency did not significantly affect blood flow and the diameter in the IVC after AVF, but markedly reduced SMC proliferation and neointimal hyperplasia, suggesting that MMP-3 is primarily involved in AVF neointimal hyperplasia but not AVF maturation. Although no proven drugs have been developed to reduce AVF failure, the strategies to inhibit SMC proliferation, ECM deposition, and neointimal hyperplasia improve AVF patency in both clinical studies and animal models.^78–80^ Moreover, emerging data suggest that endovascular AVF creation (WavelinQ and Ellipsys systems) could be an option for some patients as a minimally invasive option in addition to a sutured AVF.^81, 82^ We have shown that the mouse needle aortocaval puncture model recapitulates both endovascular AVF and sutured AVF procedures.^39, 44^ In the present study, we show that SMC-specific MMP-3 significantly reduces AVF-mediated neointimal hyperplasia of the outflow vein and the JAA, suggesting that targeting MMP-3 in SMC could be a therapeutic strategy to slow aggressive neointimal hyperplasia and improve the patency in AVF. Since AVF surgery is performed in CKD patients, it seems much better to conduct an AVF procedure in a CKD condition in mice. However, our recent study showed that the 5/6 Nx mice had much worse AVF patency than the sham mice, suggesting that CKD can facilitate AVF failure.^83^ Studies have demonstrated that MMPs are involved in the pathogenesis of CKD.^84, 85^ To avoid the effect of MMP-3 on CKD, we performed AVF in the non-CKD mice, which allow us to more specifically study the role of MMP-3 in AVF neointimal hyperplasia failure.

In summary, we show that MMP-3 is highly expressed in neointimal SMCs after AVF. Deficiency of MMP-3 in SMCs suppresses AVF neointimal hyperplasia and improves AVF patency *in vivo*. Mechanistic data show that MMP-3 regulates AVF neointimal failure likely by modulating FAK- AKT signaling. Our findings suggest that MMP-3 is a key mediator of AVF neointimal failure and that targeting MMP-3 could be a novel strategy to prevent AVF failure and improve outcomes in patients with ESKD.

## Non-standard Abbreviations and Acronyms

Arteriovenous fistulae: (AVF);
matrix metalloproteinase-3: (MMP-3);
vascular smooth muscle cells: (SMCs);
10% phosphate-buffered formalin: (10% NBF);
4% paraformaldehyde: (4% PFA);
hematoxylin and eosin: (H&E);
Verhoeff-Van Gieson: (VVG);
Dulbecco’s modified eagle’s medium: (DMEM);
glyceraldehyde-3-phosphate dehydrogenase: (GAPDH);
fetal bovine serum: (FBS);
proliferating cell nuclear antigen: (PCNA);
end-stage kidney disease: (ESKD);
tyrosine kinase focal adhesion kinase: (FAK);
Sulforhodamine-B: (SRB);
inferior vena cava: (IVC);
juxta-anastomotic area: (JAA);
tamoxifen: (TMX);
CDK: (cyclin-dependent kinase).

---

**Suppl. Fig. 1.**
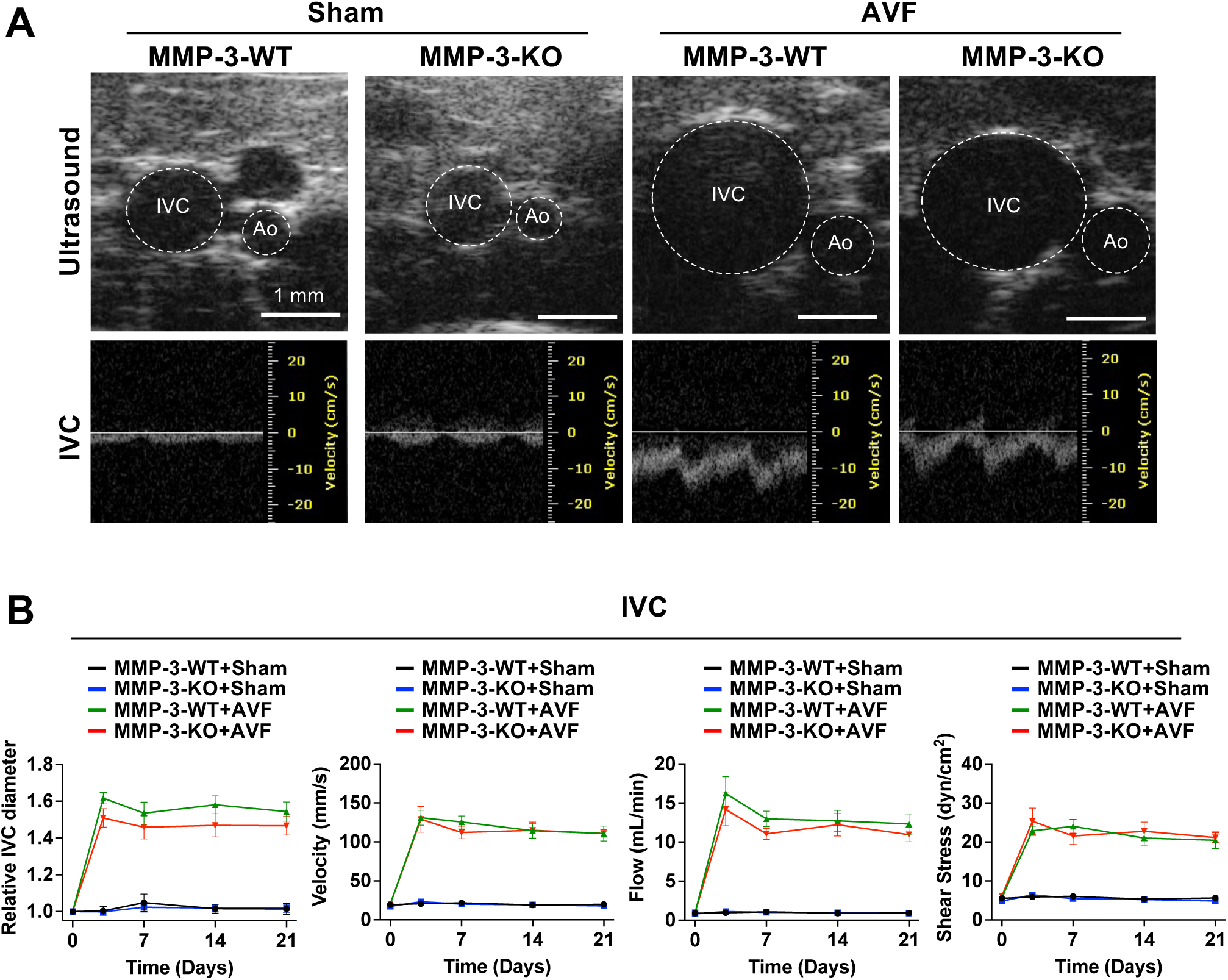
Deficiency of MMP-3 does not significantly affect hemodynamic changes after AVF. 9-week-old male MMP-3-WT and MMP-3-KO mice were subjected to AVF creation. The diameter, blood flow, and velocity of the IVC were monitored by ultrasound. Shear stress was calculated based on the velocity and diameter. **A**, Representative ultrasound showed outward remodeling in the IVC after AVF. **B**, Hemodynamic data of the IVC. Quantitative data showed that MMP-3 deletion did not significantly affect the diameter, blood flow, velocity, and shear stress in the IVC after AVF. n=6. Data were analyzed using two-way ANOVA with Tukey’s test for multiple comparisons for each time point. *P*<0.05 was significant. **AVF**, arteriovenous fistula; **IVC**, inferior vena cava; **Ao**, aorta.

**Suppl. Fig. 2.**
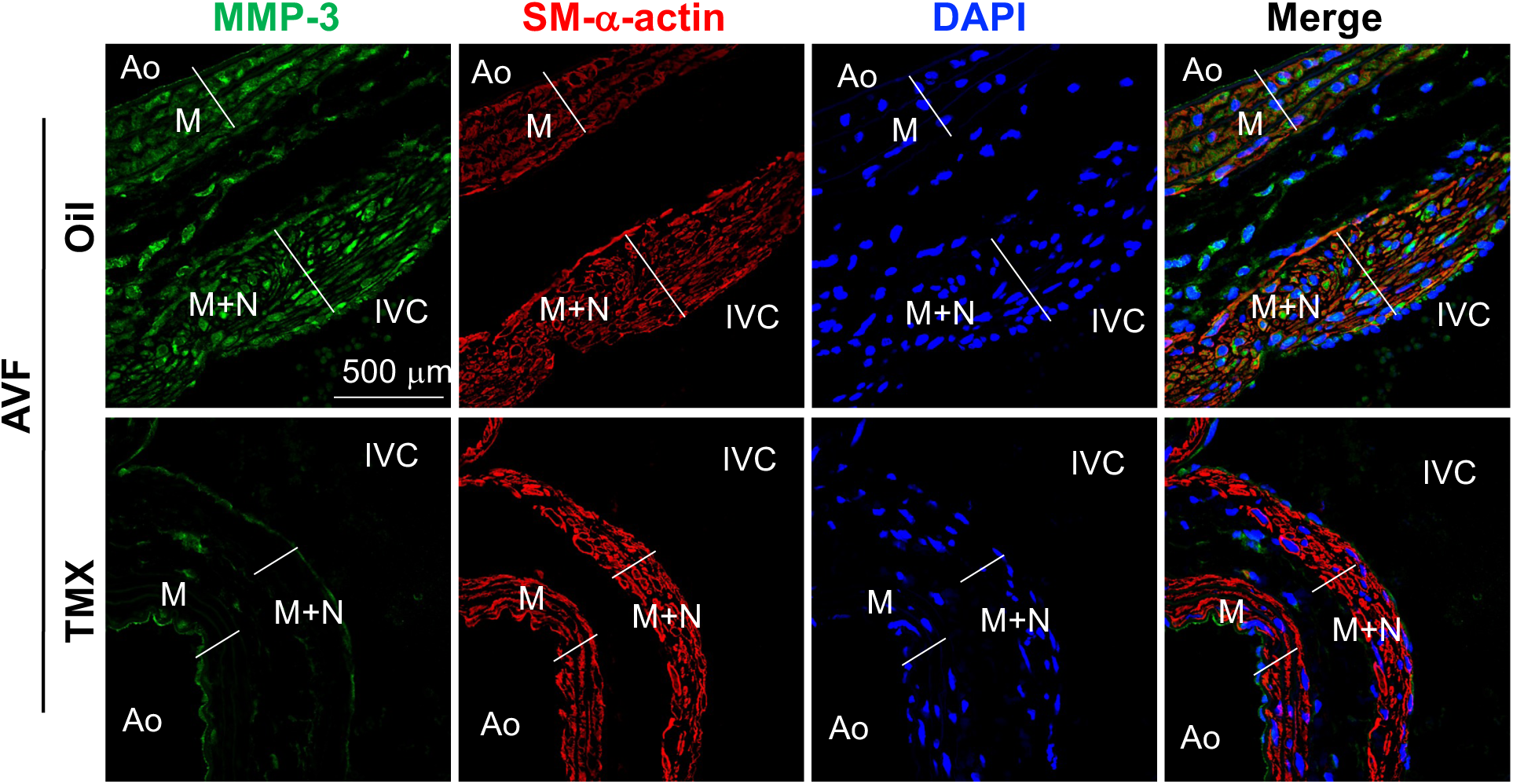
Effect of MMP-3 deficiency in SMCs on the changes in hemodynamics after AVF. 7-week-old male SMMHC-Cre^+^/MMP-3-flox^+/+^ mice were IP injected with 75 mg/kg tamoxifen (TMX) or Oil for 5 consecutive days, followed by 2 weeks of rest, and then subjected to AVF creation. Mice were harvested after 42 days. **A**, Immunofluorescence staining showed that MMP- 3 expression was largely reduced in the IVC in TMX-injected SMMHC-Cre^+^/MMP-3-flox^+/+^ mice compared with Oil-injected mice after AVF. **B**, Hemodynamic data of the IVC. Quantitative data showed that SMC-specific MMP-3 deletion did not significantly affect AVF-mediated hemodynamic changes. n=5-6. Data were analyzed using two-way ANOVA with Tukey’s test for multiple comparisons for each time point. *P*<0.05 was significant. **IVC**, inferior vena cava; **Ao**, aorta; **M**, media; **N**, neointimal area; **L**, lumen; **DAPI**, nucleus.

**Suppl. Table 1.**
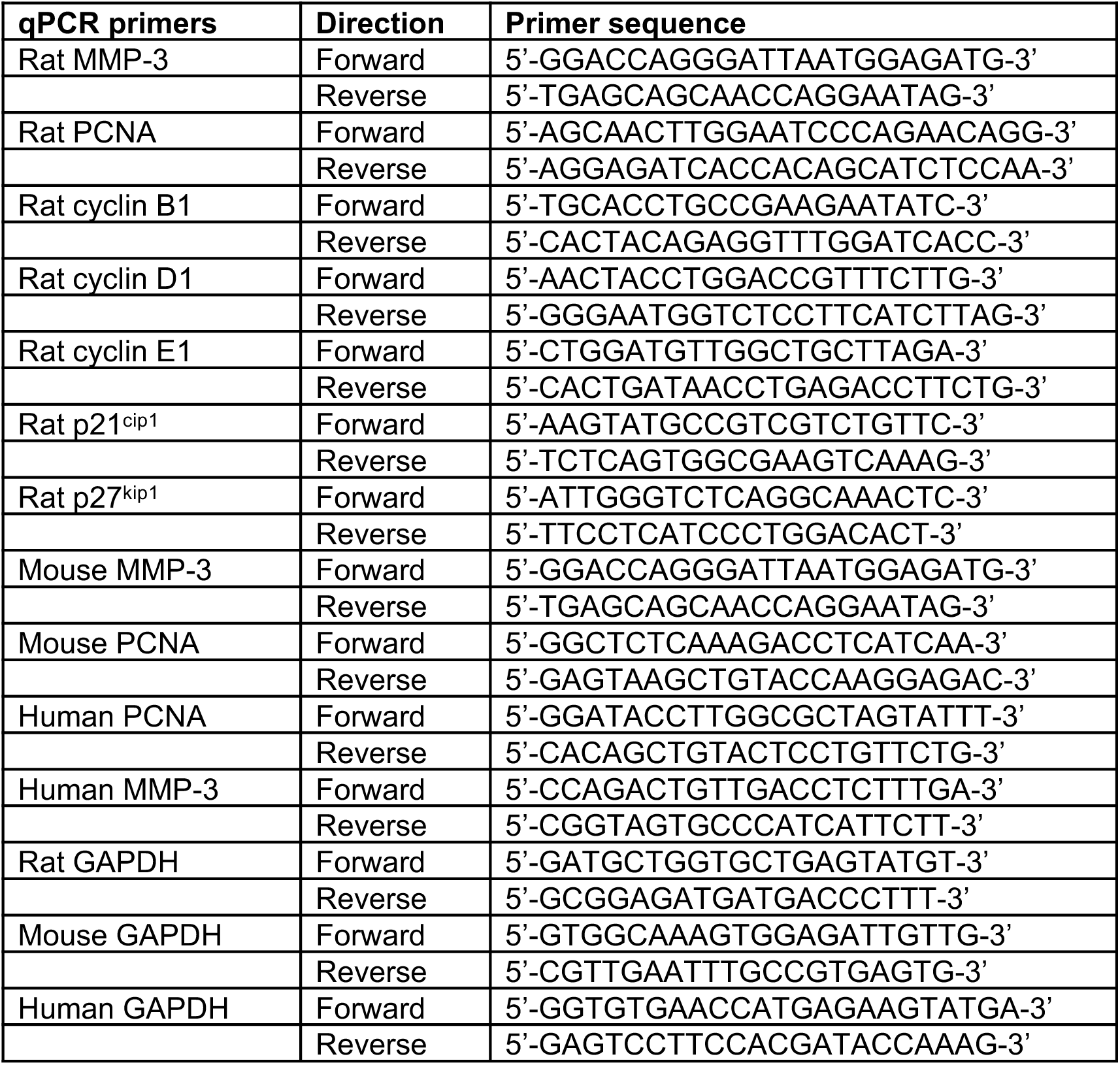

**Suppl. Table 2.**

| Target antigen | Vendor or Source | Catalog # | Working concentration |
| --- | --- | --- | --- |
| Mouse anti-GAPDH | Millipore Sigma | MAB374 | 1:10000 (WB) |
| Mouse anti-SM-α-actin | Dako | M0851 | 1:500 (IF) |
| Rabbit anti-MMP-3 | Abcam | ab53015 | 1:500 (WB); 1:50 (IF) |
| Mouse anti-Ki67 | Thermo Fisher Scientific | 14-5698-82 | 1:300 (IF) |
| Mouse anti-PCNA | Santa Cruz | sc-56 | 1:500 (IF) |
| Rabbit anti-p-AKT (Ser473) | Cell Signaling | 9271S | 1:500 (WB); 1:200 (IF) |
| Rabbit anti-AKT | Cell Signaling | 9272S | 1:500 (WB) |
| Rabbit anti-ERK 1 | Santa Cruz | SC-94 | 1:1000 (WB) |
| Rabbit anti-p-ERK1/2 | Cell Signaling | 9101S | 1:500 (WB) |
| Mouse anti-FAK | Invitrogen | 39-6500 | 1:500 (WB); 1:500 (IF) |
| Rabbit anti-p-FAK (Tyr397) | Invitrogen | 44-624G | 1:500 (WB); 1:500 (IF) |
| Mouse anti-Fibronectin | Santa Cruz | sc-271098 | 1:1000 (WB) |
| Rabbit anti-COL1A | Santa Cruz | sc-59772 | 1:1000 (WB) |

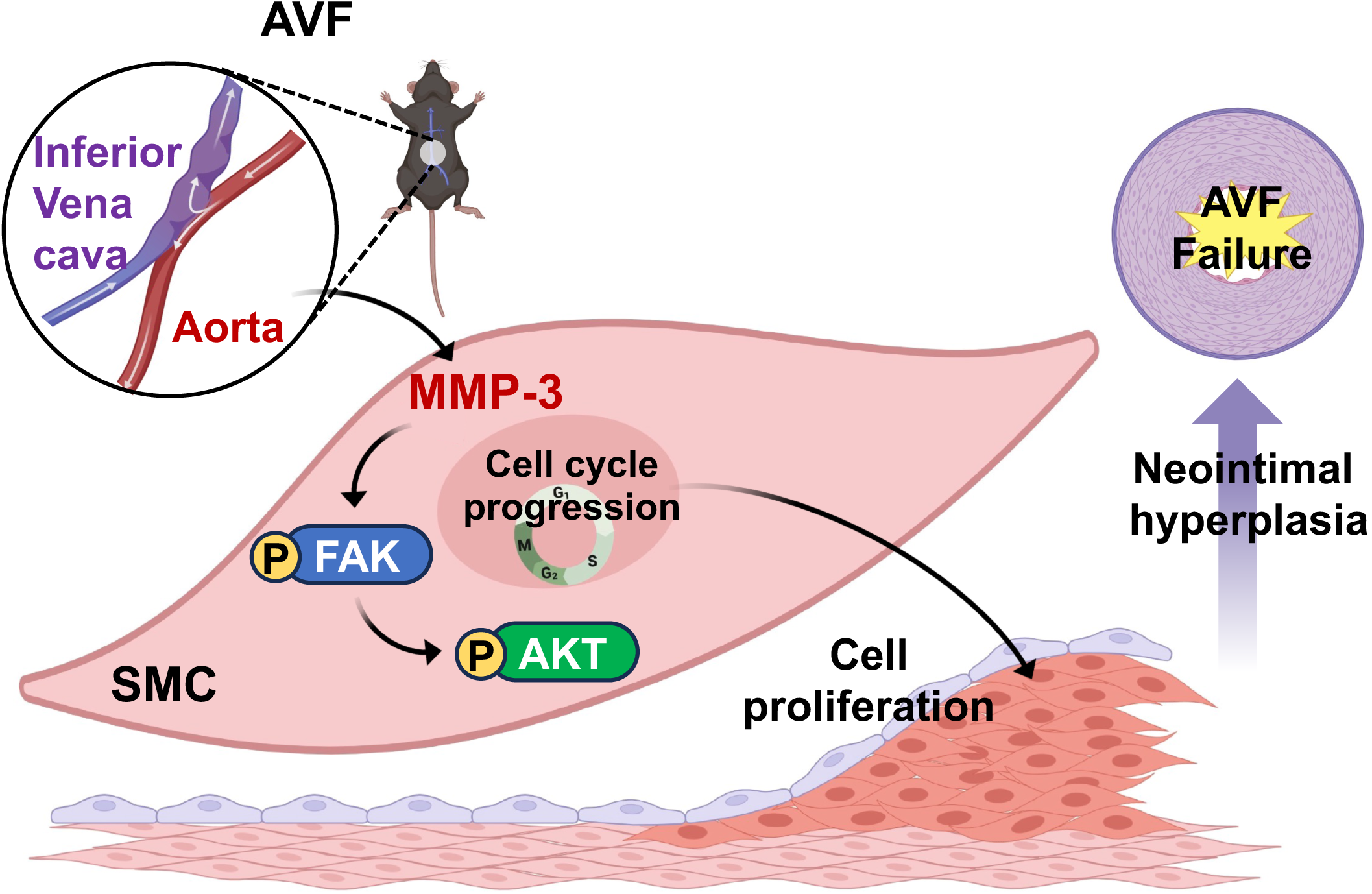
Graphic Figure.

